# Pros and cons of group averaging in studies of stochastic rodent locomotion

**DOI:** 10.1101/2025.07.17.665420

**Authors:** O. S. Idzhilova, I. S. Midzyanovskaya, V. V. Strelkov, N. L. Komarova, O. A. Chichigina

**Affiliations:** Institute of Higher Nervous Activity and Neurophysiology of Russian Academy of Sciences, 5A Butlerova St., Moscow, 117485, Russia; P. N. Lebedev Physical Institute of the Russian Academy of Sciences, Leninsky Prospekt 53, Moscow, 119991, Russia; Department of Mathematics, Univesity of California, La Jolla, San Diego, 92093, CA, USA; Physics Department, M. V. Lomonosov Moscow State University,, Moscow, 119992, Russia

**Keywords:** stochastic process, rodent, spontaneous locomotion, speed distribution, stochastic differential equation, double Rayleigh distribution

## Abstract

We analyze spontaneous walks of rodent pups in a novel environment. Individual speed distributions follow a two-component Rayleigh distribution, reflecting bimodal locomotion (progressions and lingering), while the ensemble-averaged distribution simplifies to a single exponential decay with a slower high-speed decay rate. This discrepancy arises from population heterogeneity in characteristic speeds. We propose a stochastic differential equation (SDE) for the averaged rodent, which in some cases can be derived by averaging the individual SDE. The SDE’s reveal a striking mechanistic difference in the nature of individual and averaged motion: individual rodents exhibit viscous friction (deceleration proportional to speed), whereas the ensemble behaves as if subject to dry friction (constant deceleration). We propose an algorithm of creating a “reference” rodent, which retains the shape of individual distributions but utilizes averaged scale parameters. It can be a useful personalized comparison tool to extract the information on the locomotor modes, e.g. in biomedical contexts.

## 1. Introduction

In the literature studying animal locomotion, two types of statistics are reported by different authors: some papers present probability distributions of instantaneous absolute velocity (or velocity projections, or the angle with respect to a reference line) for individual animals [1, 2, 3, 4, 5], while others work with ensemble statistics where data are collected for groups of animals and presented as a pooled set [6, 7, 8, 9, 10, 11].

The velocities of animals modeled as active Brownian particles are usually considered as a statistical ensemble. In these systems, it is even possible to determine the temperature [12]. For simple dynamics investigation, animals are often considered identical, or their characteristics of motion are averaged over many individuals, as was done, for example, for deer mice in running wheels [13]. On the other hand, animals with complex behavior and a small number of specimens are usually presented individually in the literature.

In paper [5], for example, techniques are developed to characterize the motion of an individual fly, but it is also noted that while “single fly assays are open to the criticism of being time-consuming and tedious, it should be possible to improve throughput by studying multiple arenas at the same time”, making it clear that the ultimate goal is to obtain quality group data. The main purpose of this article is to identify advantages and disadvantages of both types of statistics. We will show that neither one type is sufficient, i.e. in order to characterize animal motion in all its complexity, both individual and group statistics need to be collected, as it was done in e.g. [14, 15, 16]. On the one hand, information about individual properties is lost as a result of averaging. On the other hand, individual distributions are noisy and these random fluctuations may conceal the pattern of general properties. Even in the case of strong individual variability, some statistical characteristics, such as correlation functions, can only be obtained using data from many animals.

In this paper, we analyze tracking data from rodent pups running in an open field, see also [17]. The data include rat pups of two age groups and mouse pups. We compare statistics of motion obtained for individual rodents as well as results averaged over an ensemble of animals, which represent the movement of an averaged rodent. It turns out that the movement of the average rodent is described by a smaller number of parameters and has a simpler structure compared to that for individual rodents. In particular, the probability distribution of the instantaneous speed for individual rodents is characterized by a two-component Rayleigh distribution with characteristic speeds that describe two modes (progressions and lingering) of a cautious walk [17]. In an ensemble of animals, the distribution loses the two-component structure and also it decays slower for high speeds than the distribution for individual rodents.

We then construct a stochastic differential equation (SDE) [18, 19, 20, 21] for the “averaged rodent” speed. This equation can be derived by averaging the individual SDEs in a special case, and obtained heuristically in a more general case of the ensemble probability distribution of characteristic rodent speeds [17]. By construction, the SDE is characterized by a stationary probability distribution function that is consistent with the experimental one. This equation includes acceleration and deceleration as renewal processes [22, 23, 24] for a rodent exploring a new environment. We find that while individual rodent motion is subject to “viscous friction”, where the resistance force is proportional to the speed, averaged rodents are characterized by a constant deceleration term, which is a feature of “dry friction”.

We observe the same phenomenon in mechanics, where dry friction between two surfaces corresponds to averaging over the individual interactions of the protrusions. These individual interactions depend on the surface velocity in different ways, since the protrusions are different. As a result, the average friction does not depend on this velocity.

The similarity between these two approaches, individual and average, is that both distributions show that animals avoid moving at their average speed. This behavior observed in the context of cautious walks, and might not be typical for those mammals that are engaged in distant and energy-demanding travels, like seasonal migrations, where they move within narrow speed ranges, apparently because of gait- and speed-related optima in locomotor efficiency or biomechanical factors such as muscle and tendon stress (see [13] and references in it).

## 2. Speed distributions for rodent cautious walks

### 2.1. Speed distribution for individual animals

We used digitized data for the motion of 3 groups of animals: 13-day old (*N* = 150) rat pups (R13), 15-day old (*N* = 159) rat pups (R15), and 11-day old (*N* = 33) mouse pups (M11). These are some of the same data as were reported in our recent paper [17], see also [25]. Experimental details can also be found in Appendix A, and several typical trajectories are shown in figure A.9, demonstrating a large degree of variability among the animals. The different types of rat pup motion have been characterized by [17]. In this paper, as in [17], we focus on the “cautious walks” of the rodents, i.e., the locomotion in a new and potentially anxiogenic environment, with speeds below approximately 10-15 cm/sec. Even though higher speeds are sometimes reached by the animals in our groups, they are not maintained for long, and reliable statistics are harder to obtain, making analysis more challenging.

Figure 1 demonstrates the patterns of the absolute velocity of rat pups from group R13 by highlighting the regions of slow motion (speed *<* 1.5 cm/sec) in yellow. We can see that the speed of the animals intermittently drops to lower levels. Similar patterns are observed for other groups of rodents, as shown in figures 2 and A.10. We observe the rugged appearance of these graphs with sharp accelerations and decelerations, suggesting the presence of short-time correlations. We further note a high degree of heterogeneity among the animals with respect to the speed of motion.

**Figure 1:**
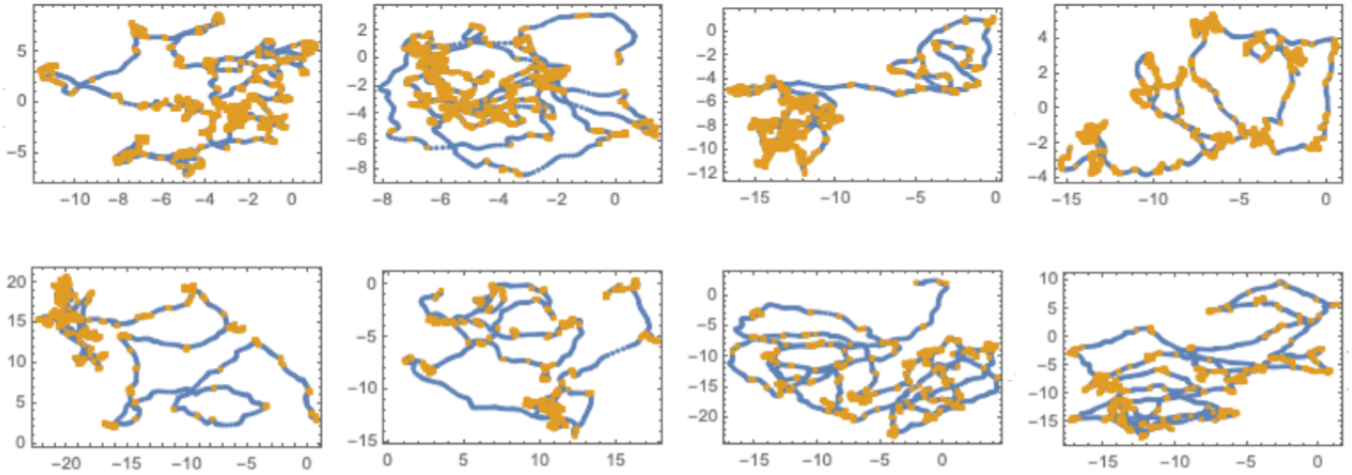
Sample trajectories of 13-day old rats, where the yellow segments correspond to speeds *<* 1.5 cm/sec, while blue segments to speeds ≥ 1.5 cm/sec. The frames show distance in cm.

**Figure 2:**
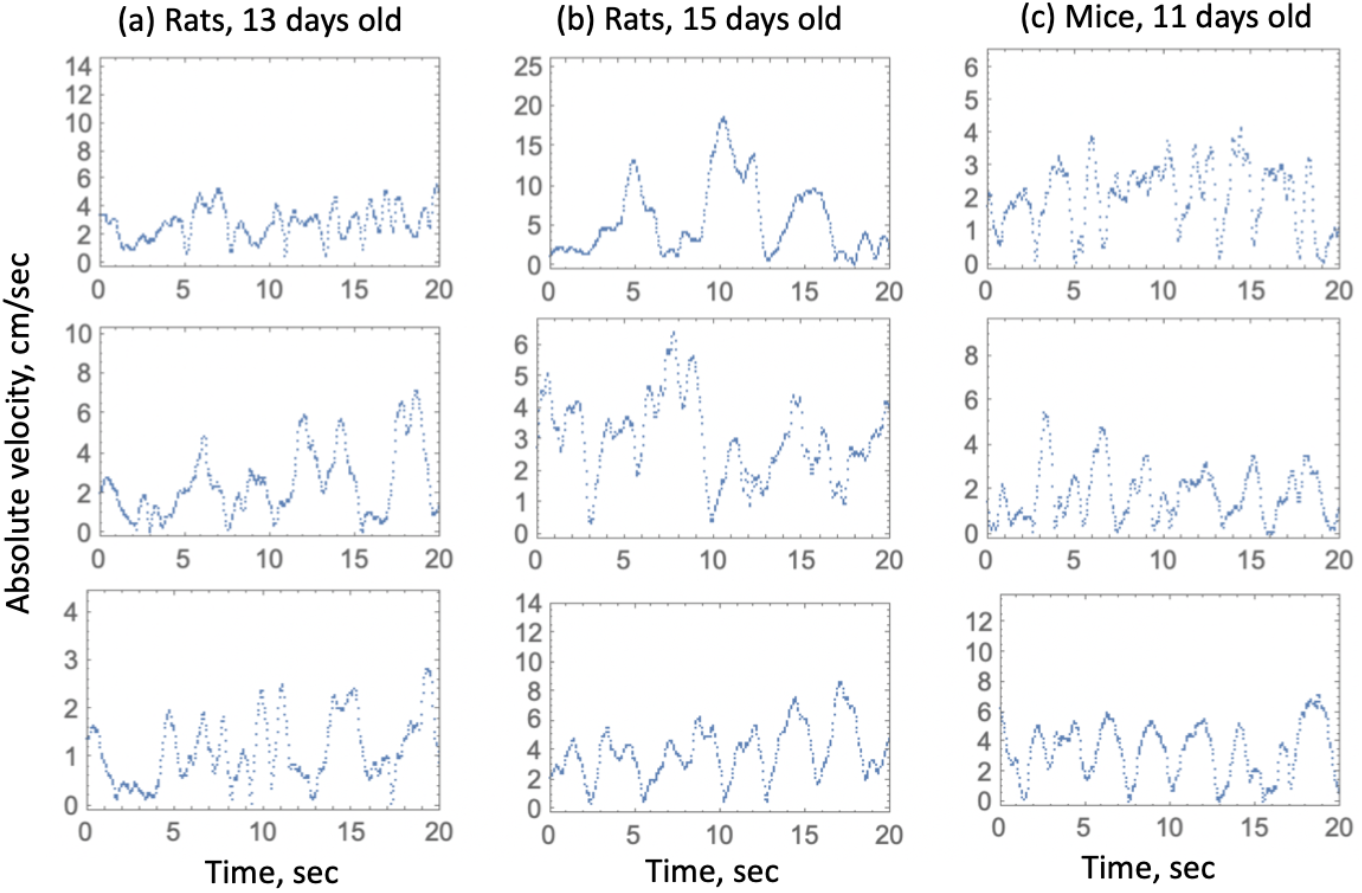
The absolute value of the instantaneous velocity, plotted against time, presented for the ffrst 20 seconds of motion. Note the difference in the vertical scale of the panels. Speed data over longer time periods are shown in figure A.10.

The process in figure 2 consists of a sequence of random pulses with the velocity frequently dropping to zero, when the rodents stop for a short time. The pulses appear to have a similar steepness of the left and right slopes, suggesting that accelerations and decelerations are of the same order of magnitude.

In [17] we studied frequency vs speed curves for individual animals in several groups: M11, R13, R15, as well as 17-day rat pups (R17), and adupt rats (Radult). These curves typically started at the origin (that is, the frequency of zero speed was zero), contained a single maximum, and decayed to zero for large speed values. We performed fitting of these individual curves with three different functions (the Rayleigh distribution, a double Rayleigh distribution, which is a weighed sum of two Rayleigh distributions, and a Gamma-distribution with the shape parameter *k* = 2). It turned out that in the majority of cases, a double Rayleigh distribution provided the best fit, which was followed by the Gamma-distribution, which was the best fit for a minority of animals.

### 2.2. Ensemble speed distribution

Here, instead of studying individual frequency vs speed curves, we pooled all the animals in each group together, and considered these curves for the whole cohort, see figure 3.

**Figure 3:**
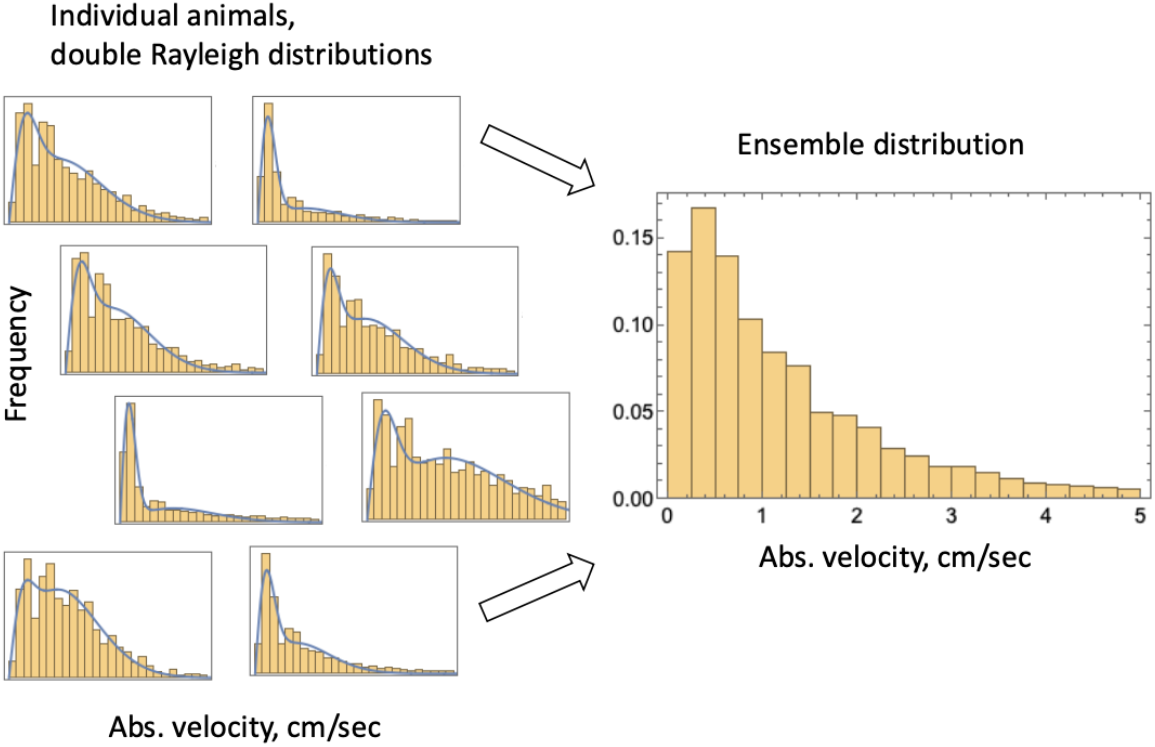
Left: Several examples of individual speed distribution for animals in group R13 (the horizontal axes spans [0, 5]cm/sec, and the blue line shows the best fitting double-Rayligh distribution). Right: the ensemble disribution of group R13.

The “ensemble” frequency vs velocity curves are presented in figure 4(a-e) for the five groups of animals, see the blue dots. These curves were fitted to several candidate functions, for each animal group:

**Figure 4:**
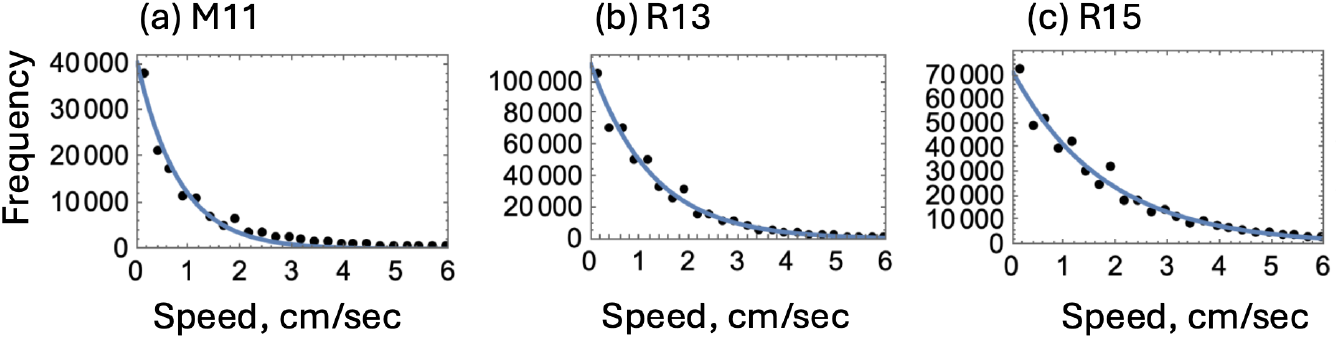
The distribution of the absolute value of the animals’ instantaneous velocity. (a-e): the frequency vs absolute velocity (over all animals at all times) for each of the groups (black dots), and the best fitting exponential function (function (i)), *f* (*v*) = *ae*^*−bv*^ (blue lines). Bine size of 0.25 was used, see figure C.11 for other examples. The best fitting values of *b* are shown in figure 5 (black circles connected with a solid line).

1. The exponential function, *f* (*v*) = *ae*^*−bv*^. When normalized, this function represents the exponential distribution.
2. A power law with the power being one of the fitted parameters, 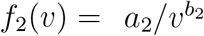, which is relevant for the Levy walks [26, 27]. This function is similar to the Pareto distribution, where it is assumed that the minimum possible value of the absolute velocity, *v*, is smaller than the minimum velocity in the dataset.
3. The function 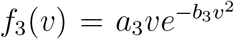, which when normalized, gives rise to the Rayleigh distribution.

In all cases, non-normalized, two-parametric functions were used, and their logarithms fitted to the logarithm of the non-normalized frequency data, by using the standard square error minimization procedure. The details of fitting are given in Appendix C, also see [17]. The best model is shown to be function (i), which, when normalized, is the exponential distribution,

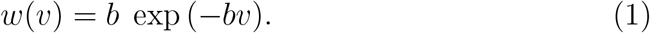

Figure 4 shows the best fitted function (i) for the three groups of animals, where the blue solid line represents the best fit. Figure 5 plots the best fitting quantity 1*/b* in function (1) for all animal groups (the black circles connected with a solid line, with the bars denoting 95% confidence intervals).

**Figure 5:**
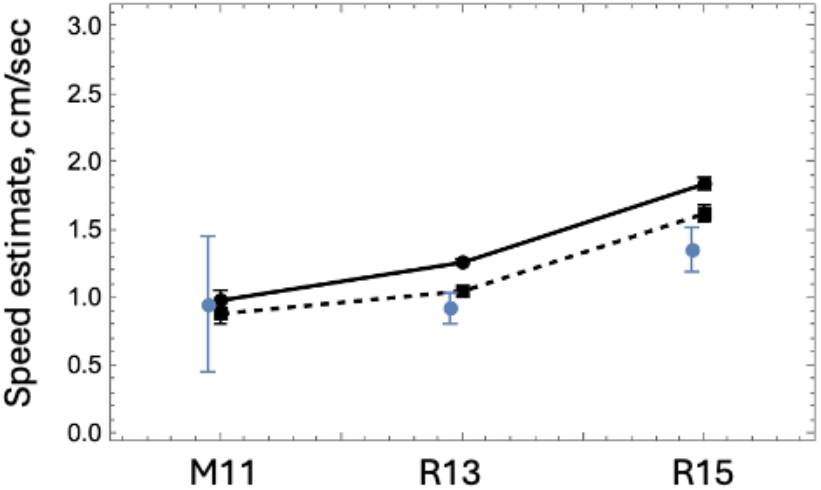
Three estimates of the characteristic speed of the ensemble of animals, for the three groups. Black circles (connected with a solid line) represent the value 1*/b* (where *b* is best fitted absolute values of the exponent in function (i), *f* (*v*) = *ae*^*−bv*^). Black squares (connected with a dashed line) represent the value *v*_Σ_ in equa<u>ti</u>on (11), fitted to the data of velocity projections. Blue dots represent the values 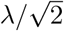 calculated by fitting the ensemble distribution of characteristic speeds with function (7) with *k* = 2 (speciffcally, function (D.2)). All fitted values are plotted with the 95% conffdence intervals represented by bars (hard to see in some cases).

### 2.3. Connection between individual and ensemble speed distributions

As it was shown in [17] in a group of animals, the speed probability distribution for a single individual *j* is described by

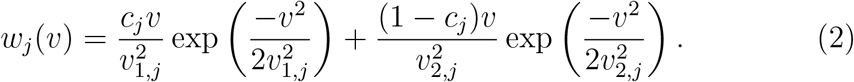

The two terms represent the two modes of cautious walks in rodent locomotion. Characteristic speeds *v*_1,*j*_ and *v*_2,*j*_ with *v*_1,*j*_ *< v*_2,*j*_ correspond to the “lingering” and “progression” modes of animal *j*, respectively. Distribution (2) can be represented through two conditional Rayleigh distributions,

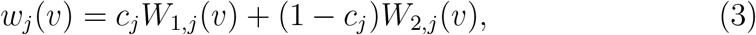

where we denoted a single Rayleigh distribution,

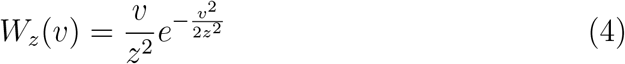

and used notation *c*_*j*_ for the weight of the lingering mode.

We pool together characteristic velocities to create a set that characterizes the whole group. In the simplest case where all the *c*_*j*_ *≈* 1*/*2, we can simply take (*V*_1_, …, *V*_*n*_, …, *V*𝒩) = (*v*_1,1_, …, *v*_1,*j*_, …, *v*_1,*N*_, *v*_2,1_, …, *v*_2,*j*_, …, *v*_2,*N*_), where 𝒩 = 2*N*. For a more general case where the two Rayleigh components come with unequal weights, the procedure of creating the common representative pool of characteristic velocities is described in Appedix Appendix D. Then the probability distribution for the velocity that characterizes a group of *N* animals is given by a group probability distribution,

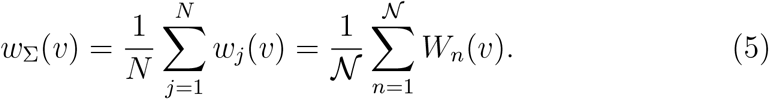

Here we will demonstrate that although the individual distributions, equation (2), decay rapidly at high values of *v*, the distribution for an ensemble, equation (5), could have a much thicker tail.

In the limit of a large number of individuals, we can use the continuous variables *V* instead of *V*_*n*_, and rewrite quantity (5) in an integral form, where (normalized) Rayleigh distributions contribute with weights defined by the representation of each characteristic velocity value in the population. This can be expressed as

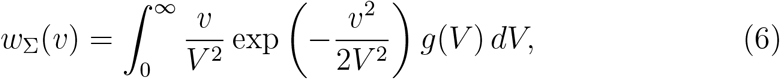

where *V* is the continuous characteristic velocity of an individual contribution 7 and *g*(*V*) is the probability distribution in the group of animals. Clearly, the behavior of the ensemble distribution *w*_Σ_(*v*) depends on the distribution of characteristic speeds, *g*(*V*), in the group of animals. This probability distribution describes the experimentally measured quantities *v*_1,*j*_ and *v*_2,*j*_ in equation (2).

To obtain the empirical probability distribution of the characteristic velocities, we used the results of [17], where we fitted distributions of absolute velocity for individual animals with function (2). The best fitting parameters *v*_1,*j*_ and *v*_2,*j*_ were pooled together as described above. In order to determine the shape of the tail of the group probability distribution of the characteristic speeds, we assumed that it had the following very general shape:

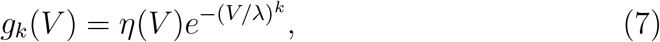

where parameter *k >* 0 defined the power of *V* in the tail of the distribution, parameter *λ >* 0 controlled the width of the distribution with a given *k, η*^*1*^(*V*) ≥ 0 for all *V*, and *η*(*V*) ≈ 1 for *V λ*, so *η*(*V*) did not affect the tail of the distribution (7). We determined that *k ≈* 2 is a good approximation (for all the animal groups parameter *k* was found to be between 1.5 and 2, see Appendix D).

Using the value *k* = 2, we can evaluate equation (6), to obtain

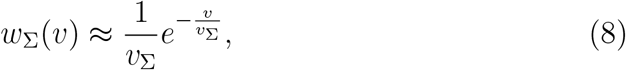

see Appendix D for the details of this calculation. We can see that, for the given empirical shape of the distribution of characteristic speeds in a rodent group, the resulting group-averaged speed distribution (8) decays exponentially, in agreement with the the experimentally-based Eq. (1) with *v*_Σ_ = 1*/b*.

This somewhat counterintuitive result can be rephrased as follows: even though individual animals contribute speeds (*v*) that are governed by Rayleigh-like distributions (with tails that decay as an exponent of a negative square of *v*), the resulting ensemble distribution has a much thicker tail (and decays as an exponent of a negative *v*). This is a consequence of the distribution, among the animals, of the individual characteristic speeds, equation (7). It is the tail of this function (parameter *k* in equation (7)) that controls the tail of the ensemble speed distribution, with the latter always decaying slower than the former. For example, the ensemble distribution in equation (8) decaying exponentially with *v*, while the distribution of characteristic speeds has *k* = 2 and exhibits Gaussian decay.

### 2.4. Ensemble velocity projection distributions

Next we turn to the projections of the instantaneous velocity on the *x*-axis. Figure 6 (top row) shows experimental distributions of the values *v*_*x*_, the *x*-projections of the animals’ instantaneous velocity.

**Figure 6:**
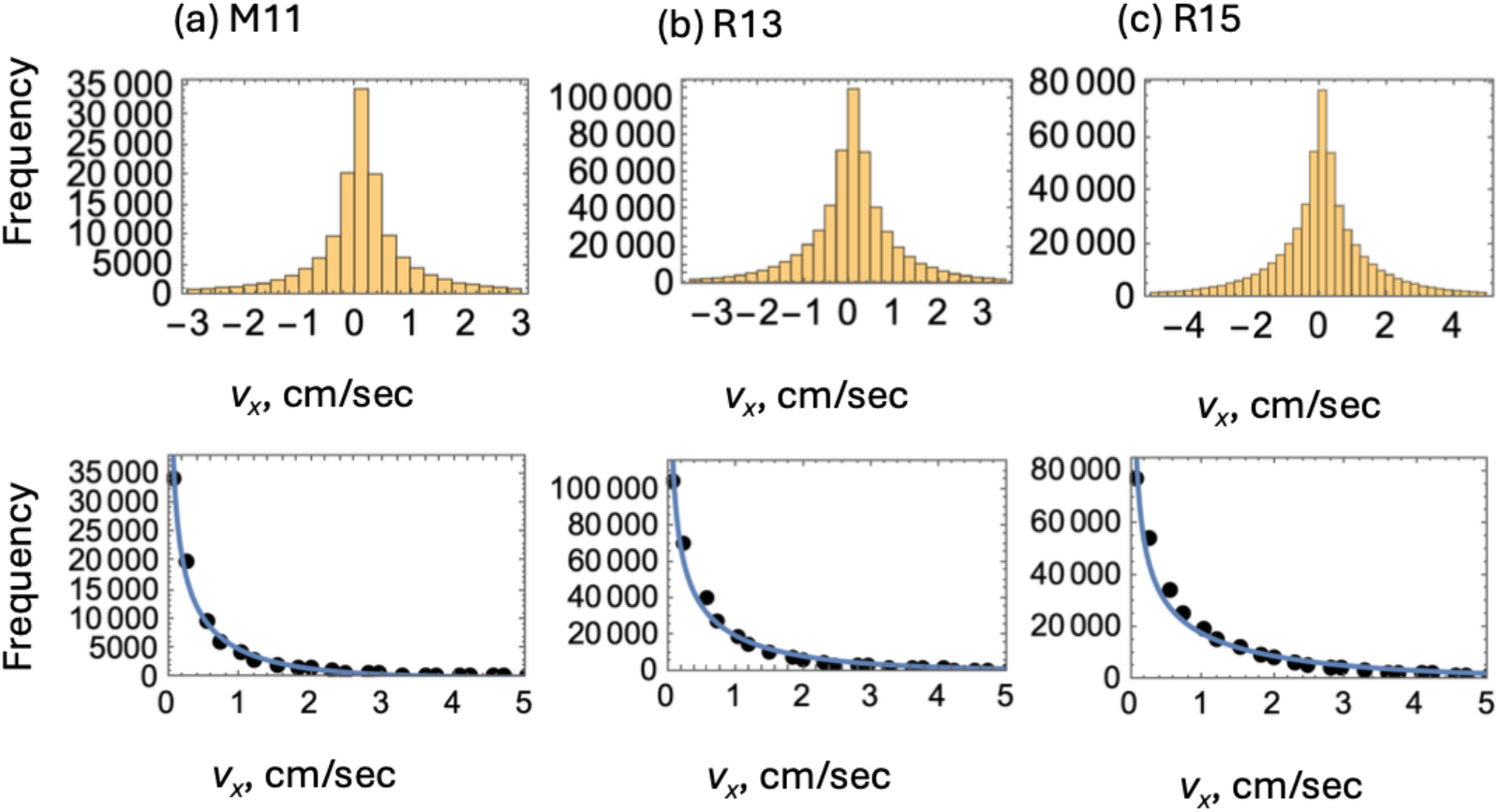
Projections of the animals’ instantaneous velocity, *v*_*x*_, for all the animal groups: (a) M11, (b) R13, (c) R15. Top row: histograms of the *x*-projection. Bottow row: the absolute value of the projections together with the best fitting function, equation (11). The best fit values of *v*_Σ_ are shown in figure 5 (black squares connected with a dashed line).

The PDF for the two projections of the velocity in two-dimensional isotropic motion is

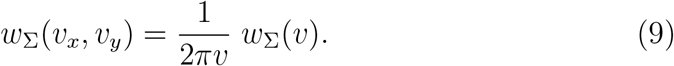

The PDF for one projection has a sharp peak at zero. This PDF can be calculated as:

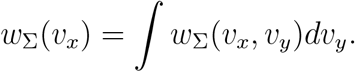

From Eqs. (8) and (9) we have:

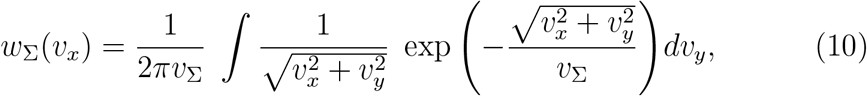

and the approximate expression is

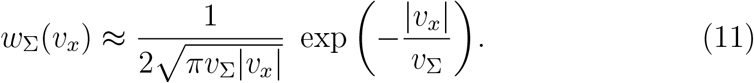

Equation (11) was used to fit the data on the *x*-projection of the velocities for the three types of animals. The best fits are shown in figure 6 (bottom row), and the best fitting values of *v*_Σ_ are plotted in figure 5 by black squares connected with a dashed line, where vertical bars represent their 95% confidence intervals. As expected, the estimates *v*_Σ_ are close to the estimates of 1*/b* obtained from fitting the ensemble distribution of instantaneous speed values.

## 3. Stochastic differential equations for rodent cautious walks

### 3.1. A general stochastic differential equation

We start with a general stochastic differential equation written as

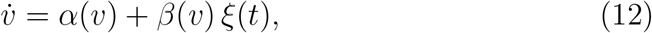

where *α*(*v*) and *β*(*v*) are functions of *v*, and *ξ*(*t*) is stationary white Gaussian noise with zero mean *ξ*(*t*) = 0 and correlation function *ξ*(*t*)*ξ*(*t* + *τ*) = 2*Dδ*(*τ*).

The corresponding Fokker–Planck equation for the velocity distribution in Stratonovich interpretation is

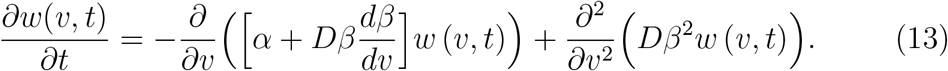

The stationary solution [28, 29, 30] is

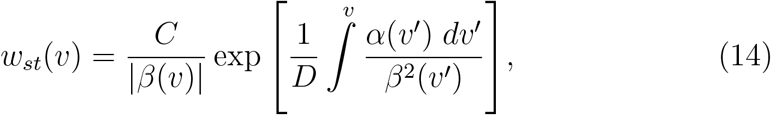

where *C* is a normalization constant.

### 3.2. Stochastic differential equation for an individual rodent speed

In [17], the change of the velocity was studied by considering acceleration and deceleration separately. We modeled the acceleration of an individual rodent as a pulse process, where each positive pulse corresponded to rapid acceleration, following a putative brain command. This process was characterized by an intensity *D* and a short correlation time. The pulses (to the first approximation) did not depend on the current velocity and were assumed to follow Poisson statistics. In the case, when several pulses arrive in a row with small time intervals between them, the rodent develops high velocities. This acceleration is compensated by a deceleration corresponding to a high energetic cost of locomotion: rodents naturally tend to decrease locomotor activity instead of moving endlessly.

The deceleration changes between weak and strong modes as a dichotomous process. This process describes switching between the locomotion and the lingering state modes. For a given rodent, during each mode the deceleration is the same as for a Brownian particle with corresponding damping coefficient *γ*_*i,j*_, where *i* = 1, 2. The deceleration is proportional to the speed, since the higher the speed, the more effort is required. As a result, PDF decreases quickly at large velocities and corresponds to a normal distribution for the projection of speed and to Relay distribution for the absolute value of the speed.

The correlation time of switching decelerations obtained in [17] is much longer than the correlation time of acceleration. The short correlated acceleration is represented as white noise. The corresponding Langevin equation for two-dimensional motion is presented in Eq. (20) [17]:

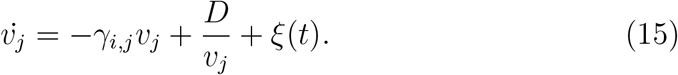

The first two terms correspond to the deterministic part *α*(*v*) in equation (12) and *β*(*v*) *≡* 1. The corresponding Fokker–Planck equation is

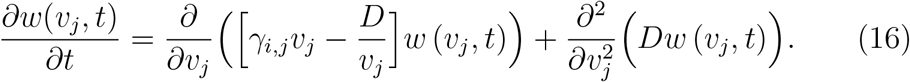

For rodent *j*, the stationary distribution during mode *i* is obtained by substituting them in (14):

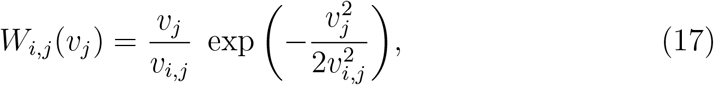

where 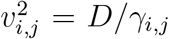. This Rayleigh distribution has the following moments: 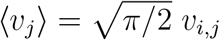, and 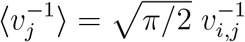.

### 3.3. Stochastic differential equation for the speed of an averaged rodent

Let the ensemble of *N* individual rodents with the corresponding two damping coefficients be used to investigate the properties of the averaged rodent. We effectively replace a population of *N* rodents with two characteristic speeds each, with a population of 𝒩 rodents with a single characteristic speed, which we denote *V*_*n*_, *n* = 1,…, 𝒩. The speed of each such rodent *n* is distributed according to

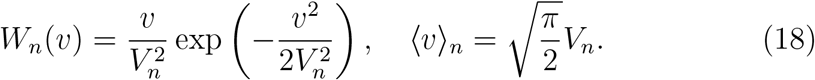

We have 𝒩 different damping coefficients (*γ*_*n*_) and SDEs for each of the individual rodents. To find SDE for the averaged rodent’s velocity *v* = *(v*_*n*_), we sum 𝒩 equations for the individual rodents, equation (15).

Assume that the distribution of the parameter *V*_*n*_ among the rodents is given by

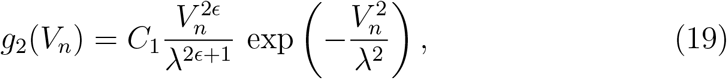

where *C*_1_ is a normalization constant, and *λ* and *ϵ* are constants responsible for different aspects of the behavior of this distribution, as explained below. The fact that the tail of the distribution has a Gaussian decay is motivated by finding the experimentally observed distribution of characteristic speeds, see equation (7) with *k* ≈ 2, also Appendix D. In the limit *ϵ* → 0, function (19) corresponds to the half-normal distribution. In the special case where *ϵ* = 1, we obtain the Maxwell distribution. For nonzero but small values of *ϵ*, it provides an adequate description of the experimentally observed distribution, figure D.13. In particular, it matches the distribution that was obtained from the experimental data, *g*_*k*_(*V*), equation (7) with *k* = 2, if we set 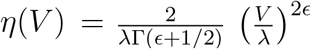, see Appendix D.

To develop the procedure where we transit from individual rodents to an averaged rodent, it is instructive to start with the special case of *ϵ* = 1, because several results admit simple analytical expressions. We then consider the more general and biologically realistic case of small ϵ.

#### 3.3.1. Maxwell distribution of the characteristic speeds

Taking ϵ = ϵ_*M*_ = 1 in expression (19), we can see that

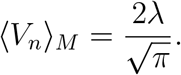

The probability distribution for the speed of the averaged rodent is given by

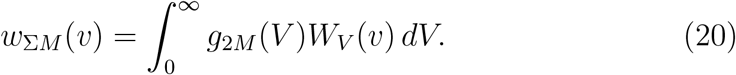

Using (19) and (18), we obtain

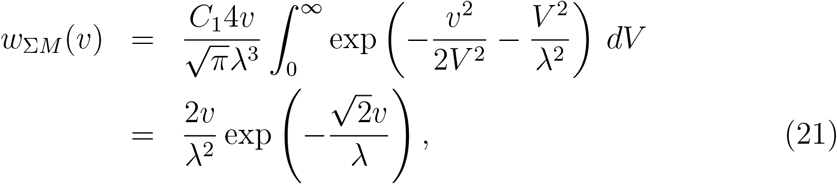

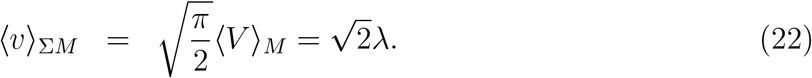

Turning to the stochastic differential equation for the motion of an averaged rodent, we note that in this case, we do not attempt to derive it from some known laws of locomotion. Instead, we use the speed probability distribution (equation (21) in this case), and choose the SDE that is compatible with this, with the overall goal to conjecture the laws that govern the animal locomotive behavior. Note that clearly, the choice of the SDE is not unique, given that the expression for speed probability distribution, equation (14), only depends on the ratio of its deterministic and stochastic components. Using the simplest equation we postulate

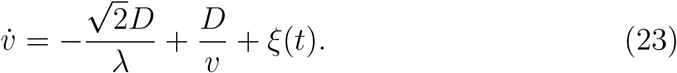

This equation is characterized by a stationary solution (14), and in addition it resembles the equation for individual rodents (equation (15)) in that the second term on the right hand side is identical, and the ffrst term describes deceleration and can be viewed as an averaging of the ffrst term in equation (15).

Next, we will show that an averaging procedure indeed gives the ffrst term in equation (23). We note that the first term in equation (15), which is first averaged over the speed values for an individual animal and then over the characteristic speeds for the ensemble, becomes

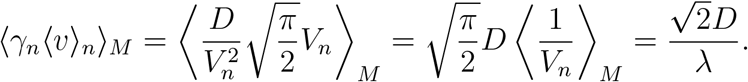

The corresponding Fokker–Planck equation is

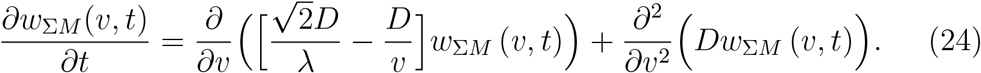

This corresponds to our earlier assumption that the individual rodents’ characteristics are encoded in the deceleration coefficient (*γ*_*n*_), while acceleration and therefore noise terms are common for all animals.

#### 3.3.2. A more realistic distribution of the characteristic speeds

The major drawback of model (19) with *ϵ* = 1 is the weak description of low speeds in figure D.13. Therefore we will next consider distribution (19) with *ϵ «* 1. Again, using equation (20), we obtain

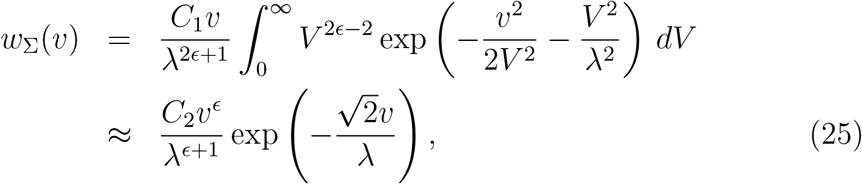

where *C*_2_ is a normalization constant and we used the fact that *V* ^2ϵ*−*2^ changes slower than the exponent. This distribution becomes exponential as *ϵ →* 0. As before, there are many ways to write an SDE to match a stationary distribution. Here we use equation (23) as a basis and modify the second term (that is responsible for the behavior near speed zero), to obtain

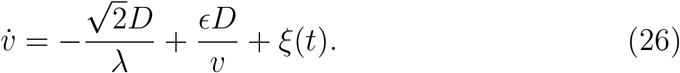

Using equation (14) it is easy to see that this SDE results in the probability distribution for the speed of an averaged rodent, *w*_Σ_, given by (25).

The corresponding Fokker–Planck equation is

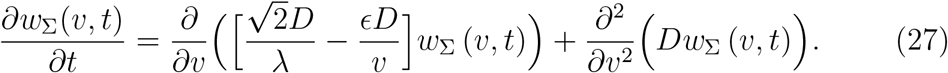

Probability distribution of the averaged velocity projection *w*_Σ_(*v*_*x*_) can be found using the same method of averaging as in Eq. (25) with normal distribution for individual projection. It is the same as in Eq. (11).

To summarize, in our model of the averaged rodent, equation (26), the nontrivial dependence on individual parameters, especially on *v*_*n*_, disappears in the process of averaging. A s a result, we replace this part of the equation by a constant deceleration 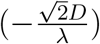. Deceleration of an averaged rodent occurs when the it is moving with *v >* 0, and the second term in the equation prevents *v* from becoming zero or negative.

The speed of the averaged rodent has a PDF slowly decreasing as an exponential function. The mechanism of high speeds damping is weaker than that of an individual animal. This distribution has a longer tail than the usual PDF corresponding to the Brownian particle model. Both models (individual and average) have a negative component in the deterministic part of SDE, but while the individual rodent’s “friction” depends linearly on *v* (15), the deceleration of an averaged rodent is constant (26). In the latter case, relatively high velocities are more probable than in the former.

Note that this methodology gives an alternative way to calculate the characteristic speed of an averaged rodent. From equation (25) we canse that the decay rate of the ensemble distribution of speeds is given by 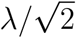, where *λ* is the parameter in equation (7) with *k* = 2 (see also equation (19)). We fitted this distribution to the empirical data (Appendix D), an plotted the best fitting values of 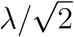 in figure 5, see blue symbols with error bars representing 95% conffdence intervals. One can see that these values are close to the characteristic speed values measured directly from the ensemble distribution of instantaneous velocity (black squares, solid lines) and its projection (black squares, dashed lines). This demonstrates self-consistency of our theory.

### 3.4. Discussion and Conclusions

The analysis of rodent locomotion presented in this study reveals fundamental differences between individual and ensemble-averaged motion statistics. At the individual level, rodent speed distributions exhibit a two-component Rayleigh structure, reflecting the bimodal nature of cautious walks, characterized by alternating phases of progression and lingering. The two-component structure disappears upon ensemble averaging, where the distribution is simplified to a function with an exponential decay. The origin of this qualitative difference between the individual and the ensemble-averaged distributions arises from the heterogeneity in characteristic speeds across the population, as described by the distribution *g*(*V*) in Equation (6).

Based on these observations we would like to propose a distinction between an *averaged rodent* and a *reference rodent*. An averaged rodent has speed statistics that is obtained simply by combining all the individual instantaneous speeds, see figure 4. This operation leads to a qualitative change in the nature of the distribution (from a double-Rayleigh distribution for individual rodents to an exponentially decaying distribution). On the other hand, a reference rodent retains the same shape of the distribution as individual rodents, but is characterized by scale parameters that are averages of the individual ones.

The algorithm of construction the reference distribution is as follows. Starting with individual distributions, find the most suitable mathematical expression (that contains parameters of shape and scale) for each of those distributions and fit the parameters. The goal is to find a description where scale parameters may be different for individual animals, but shape parameters are the same for all. Using this distribution, average all the scale parameters over the ensemble of animals. The resulting reference distribution will retain the shape of individual distributions and feature averaged scale parameters. We propose that the characteristics of this reference distribution are the most useful generalization of individual experiments, which is suitable for biological interpretation and comparison with other groups of animals. Speed distributions for reference rodents based on our data are shown in figure 7. Unlike the averaged distributions (figure 4) they have a double-Rayleigh shape with two characteristic speeds.

**Figure 7:**
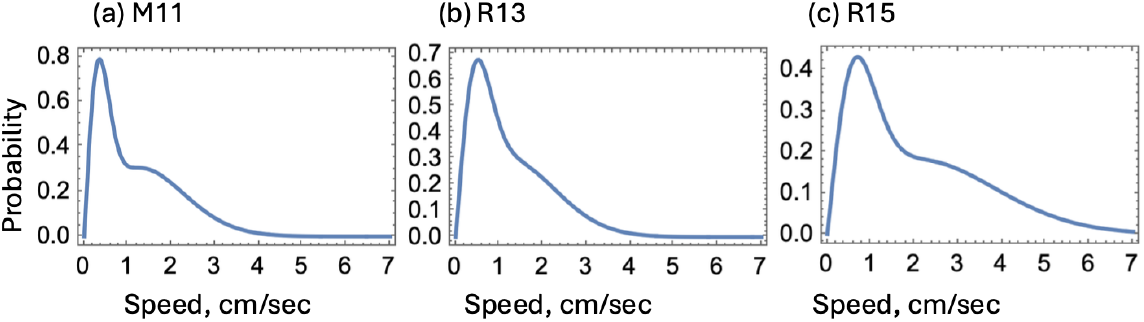
Reference rodent speed distributions constructed by averaging the individual parameters 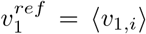 and 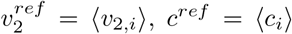 over ensembles of animals and using the double-Rayleigh form, 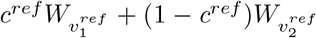 (see equations (3,4)). The characteristic speed parameters are: (a) For 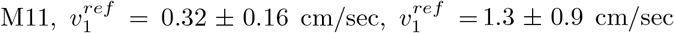; (b) for 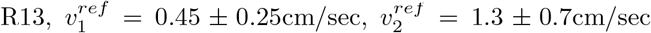; (c) for 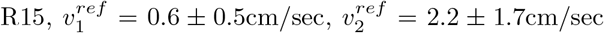 (here both mean and standard deviation are listed).

We propose SDEs both for the speed of an individual rodent and for the speed of the averaged rodent. These SDEs lead to the speed distributions that agree with the experimental ones. The individual rodents SDE assumes viscous friction, where deceleration is proportional to the speed, akin to damping of a Brownian particle. In contrast, ensemble-averaged motion is better described by a constant deceleration term, reminiscent of dry friction in macroscopic systems. This shift in dynamics mirrors physical systems where microscopic interactions (e.g., viscous friction between surface protrusions, see figure 8) average out, to yield a macroscopic dry-friction law. The SDE for the reference rodent is the same one as constructed for individual rodents, but using the averaged parameters. It provides the experimenters a possibility to quantitatively study the speed parameters of the fractal (???) intermittent locomotor modes, which is hard to do otherwise.

**Figure 8:**
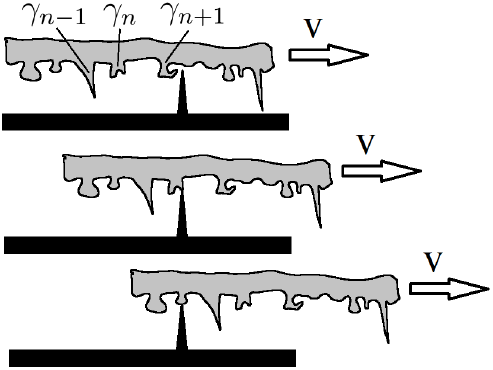
The dry friction model as an averaging over individual interactions with the moving surface’s protrusions. Protrusions of varying shapes and sizes on the upper surface are characterized by different resistance coefficients, *γ*_*n*_. Each individual interaction between these protrusions and a given protrusion on the lower surface contributes to the total braking effect of friction between the surfaces in a velocity-dependent manner. These contributions are proportional to the velocity but with different coefficients. At the microscopic level, the velocity fluctuates, corresponding to the jittery motion of moving over bumps. As a result of summing these random contributions from individual collisions between surface protrusions, the velocity dependence disappears.

The paradoxical finding that ensemble averaging produces a less Gaussian-like distribution than individual data, at first glance seems to contradict the common understanding of the Central Limit Theorem. Note that the exponential (or quasi-exponential) ensemble distribution arises not from interactions between rodents (which are absent in our experiments) but from the distribution of characteristic speeds across the population. These results emphasize the need to account for ensemble structure when interpreting aggregated data.

Our individual distribution is, in fact, a conditional distribution, given the observed values of the characteristic speeds. The variation in these characteristic speeds reflects the intrinsic properties of the physical reality we are studying, rather than being merely a consequence of measurement inaccuracies of some well-defined, unique quantity. In this regard, the problem bears greater resemblance to quantum mechanics than to classical mechanics. In our case, the diversity in characteristic speed values cannot be eliminated by increasing the number of measurements or through averaging. In a sense, the opposite is true: the uncertainty in this characteristic velocity introduces additional uncertainty into the statistical analysis of individual speed distributions and may even distort qualitative features of the distribution. It is possible that the characteristics speed *v*_1_ of one animal is equal to *v*_2_ of another, thus blurring the shape of the distribution and making it impossible to distinguish the two modes in the averaged distribution. Thus, there is no genuine violation of the Central Limit Theorem.

In conclusion, the choice between individual and ensemble analyses should be guided by the scientific question at hand. Individual statistics are essential for understanding behavioral diversity and mechanistic details, while ensemble averages provide a tractable description of group-level phenomena, which could be useful for testing hypotheses about parameter distributions. The similarity of the individual and average distributions lies in a higher coefficient of variation than that of a Brownian particle. This randomness makes the rodent movement unpredictable and thus difficult for a predator to detect and catch. The reference animal model that we construct here combines the advantages of individual and ensemble approaches. It retains the functional shape of the individual speed distributions but utilizes scale parameters that are ensemble averages. In order to understand the qualitative nature of the movement, one could use reference rodents for comparison with individual animals in the same group, or compare reference rodents among different groups. Future studies could extend this framework to other locomotor parameters (e.g., turning angles or acceleration bursts) or explore its applicability in ecological and biomedical contexts, such as collective motion or personalized therapeutic interventions.

## 4. Author contributions

Conceptualization I.M., V.S., N.K., and O.C. Data curation O.I. and I.S. Formal analysis O.C., V.S., and N.K. Methodology I.M., V.S., O.C., and N.K. Project administration V.S. Resources I.M. Software N.K. Writing – original draft N.K., O.C., I.M., and V.S. Writing – review and editing all authors.

## Appendix A. Experimental methodology

Here we provide some details of the experimental data used in this paper, which were also reported recently in [17]. Laboratory bred rodents (rats and mice) were used in this study. The animals in groups R13, R15, R17, and M11 were of both sexes. Adult rats were 4-5 month old males, with body weights of approximately 250-400 g. All the digitized tracks had length between 120 and 600 sec for all the animals. All the data can be fount at [25].

Groups M11 and R13 were both pre-visual, 2-3 days prior to eye opening. Cohort R15 had started to open their eyelids, but had not yet acquired visual sensing [31], and cohort R17 was able to fully possess the visual sensory inflow [31].

The testings were performed with no environmental adaptation period. The rodent nestlings were intact until the day of the experiment. On the test day, the mothers were gently separated from their litters, and the pups were allowed to calm down for about 30 minutes. Then, each pup was gently grasped from the huddle periphery and transported individually to the experimental arena next door. Each session started by the rodent placement to the arena center, where the animal motion was recoded. The arena was cleaned after each individual session.

Movement tracking was done offline by using tracker software ToxTrack, the algorithm ToxID [32]. This algorithm determined the body position by calculating the white mass center against the black background, every 0.04 seconds.

The experiments were carried out in accordance with the National Institute’s of Health Guide for the Care and Use of Laboratory Animals; the experimental protocol was approved by the Institutional Animal Care Committee. All efforts were made to minimize animal discomfort.

## Appendix B. Additional examples of rodent trajectories and instantaneous speed vs time graphs

Typical trajectories are shown in figure A.9, three for each type of animal in groups R13 (a), R15 (b), and M11 (c). We note the presence of a large degree of variability in trajectory patterns, with some animals remaining within a few cm^2^ of their starting position, while others traversing the arena several times, and yet others spending time within a small area as well as engaging in longer runs.

**Figure A.9:**
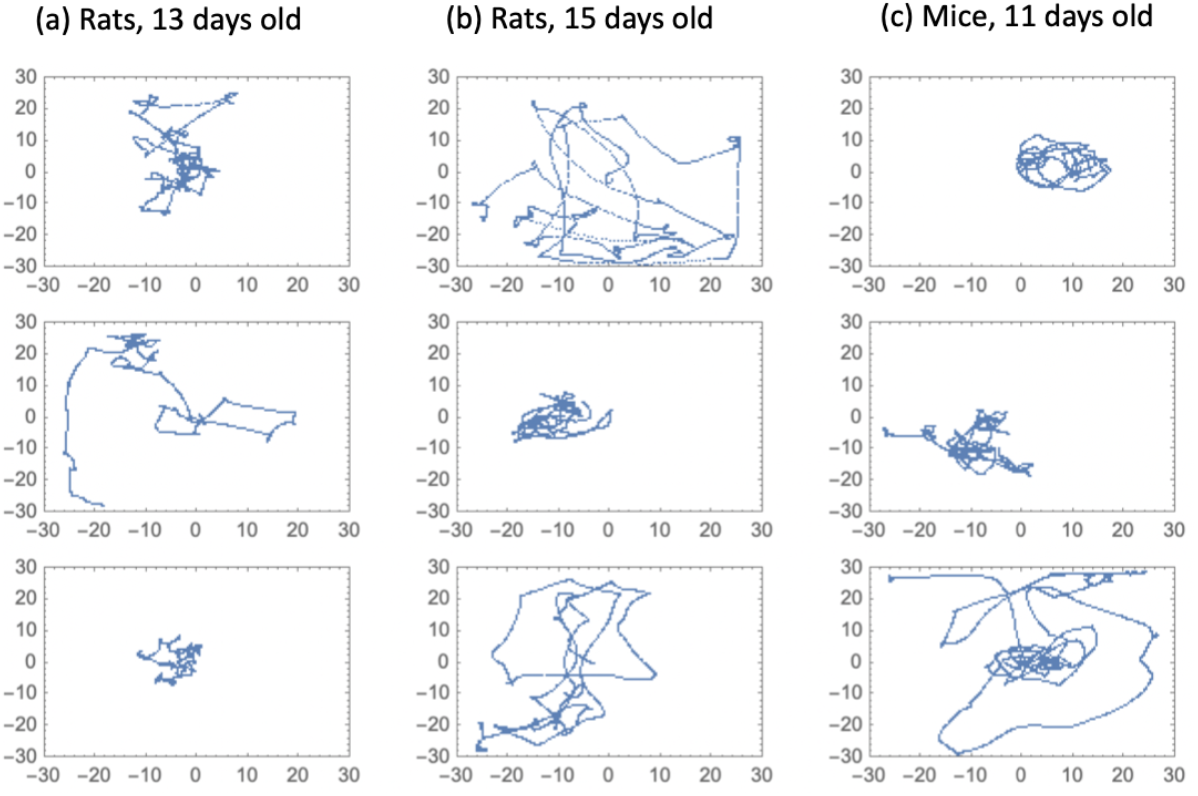
Trajectories corresponding to blind walks of (a) 13-day old rats, (b) 15-day old rats, and (c) 11-day old mice. The frames show distance in cm.

Figure A.10 plots the the instantaneous speed of animals as a function of time, with figure 2 zooming into a shorter interval of motion.

## Appendix C. Fitting the frequency vs absolute velocity graphs for all animals in a group

The fitting procedure used for this system was described in detail in [17]; in particular, while the usual maximum likelihood algorithm shifts the emphasis toward the tails of he distribution, a bin-based method captures the relevant regions of cautious walk speeds, and is relatively stable with respect to changes in the bin-size. Therefore, here we continue to use the bin-based method, as described below.

We perform a comparison of fits of the frequency vs absolute velocity graphs (over all the animals in the groups) with functions (i)-(iii) in Section 2.2. In figure C.11, all three functions (with the best fit parameters) are plotted together for comparison. The columns of the figure correspond to the three animal groups and the different rows to different bin sizes.

To compare the three functions, we have calculated the *R*^2^ value for the fits. Since all three functions have the same number of parameters, a larger value of R2 implies a more successful model. This is presented in figure C.12. It indicates that the exponential function (i) gives the best fit for all the animal groups and for all the bin sizes.

**Figure A.10:**
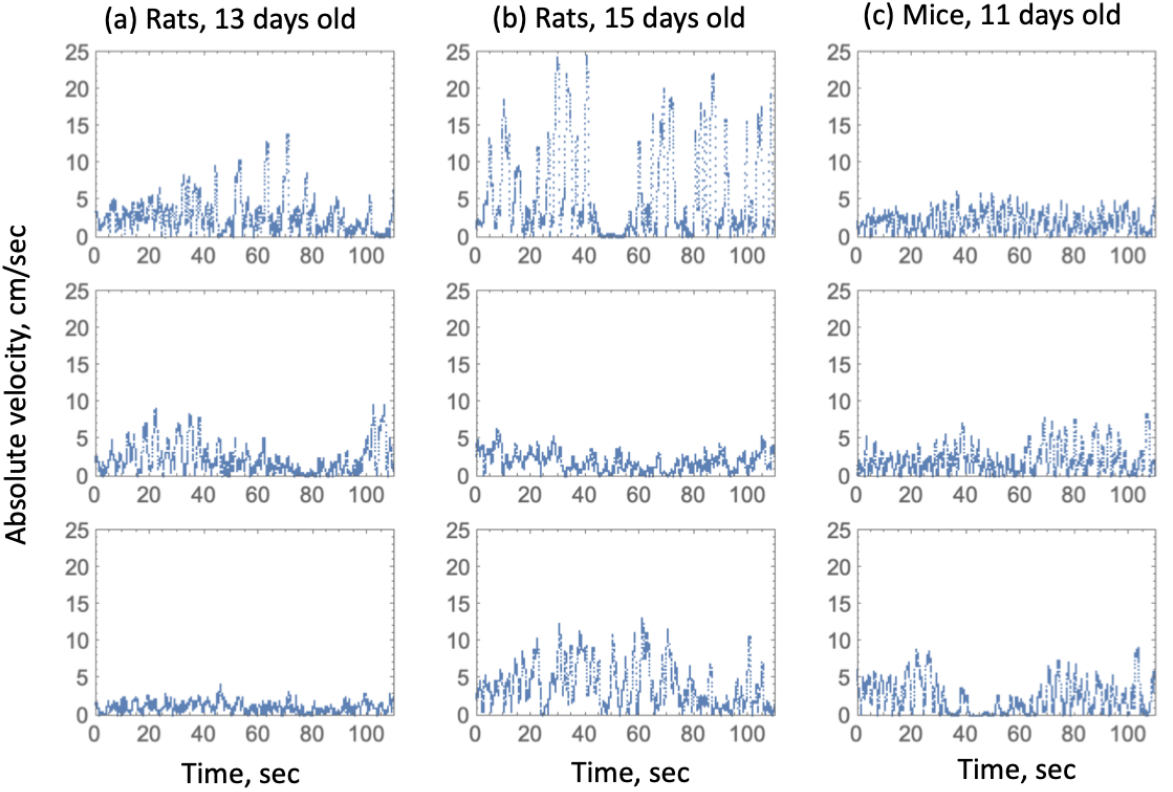
The absolute value of the instantaneous velocity, plotted against time, for the same animals whose trajectories appear in figure A.9. (a) 13-day old rats, (b) 15-day old rats, and (c) 11-day old mice.

## Appendix D. Finding *k* in Eq. (7)

In order to create an empirical set of characteristic speeds in an ensemble of animals, which takes account of non-equal weights in the two-mode distribution (3), we adopted the following procedure. We fixed an integer *M* = 10 (the higher this number, the more precisely the weights are taken into account). For each animal *j* in a given group, we calculated *n*_1_ = [*c*_*j*_*M*] (where the square brackets denote rounding) and *n*_2_ = *M − n*_1_. We then included *n*_1_ “copies” of the value *v*_1,*j*_ and *n*_2_ “copies” of the value *v*_2,*j*_. As a result we obtained a set of 𝒩 = *MN* characteristic speed values. These sets for the three animal groups are represented as yellow histograms in figure D.13. They have been normalized to represent probability distributions.

In order to extract the value *k* in the exponent of function (7), we used a specific example of this class of functions, the Weibull distribution:

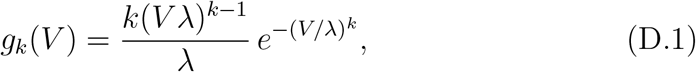

where parameter *k >* 0 deffnes the power of *V* in the tail of the distribution, and parameter *λ>* 0 controls the width of the distribution with a given *k*.

**Figure C.11:**
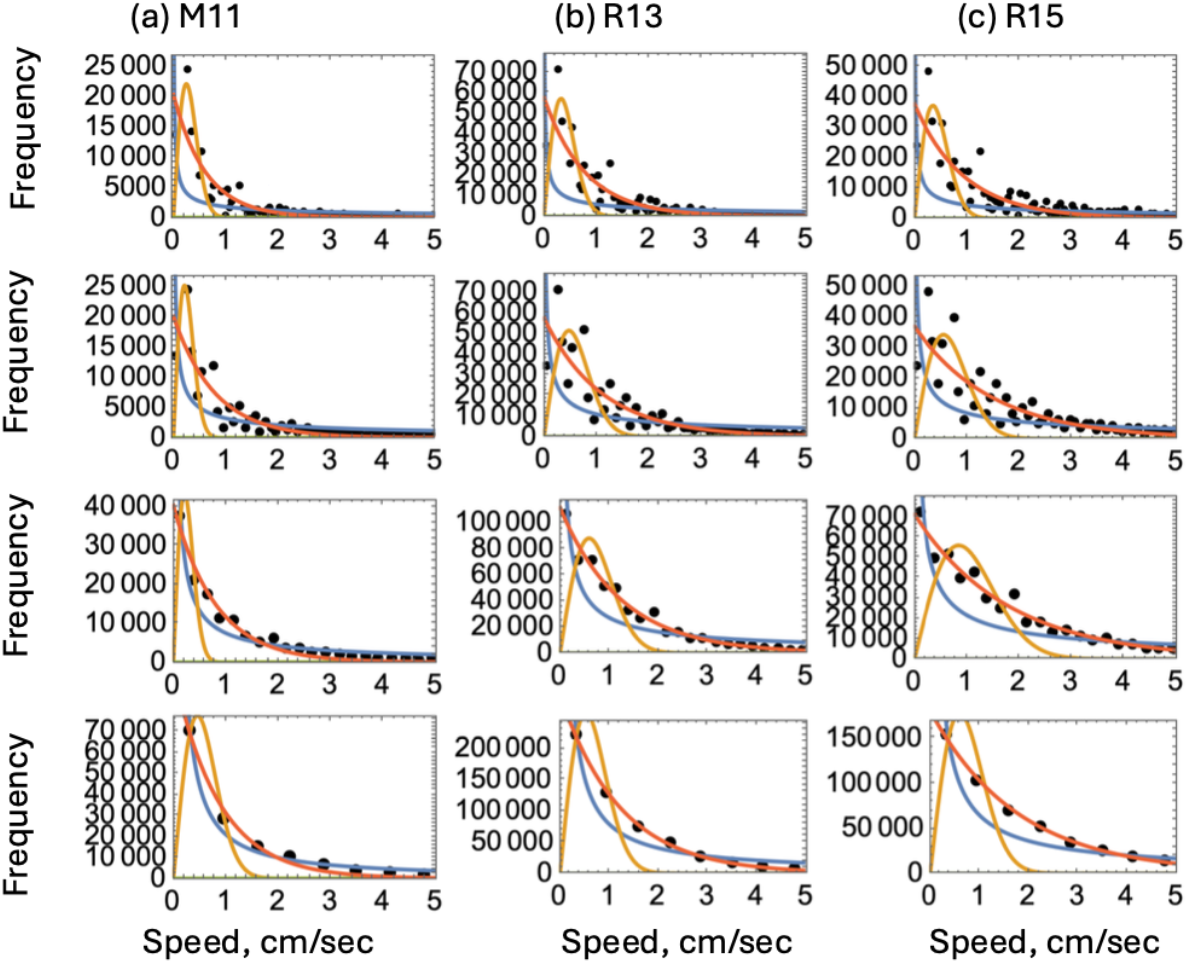
Frequency of the absolute value of the instantaneous velocity for ensembles of animals (black dots), plotted together with the best fits of the three functions: (i) (exponential, red), (ii) (power-law, yellow), and (iii) (Rayleigh, blue), for the three animal groups. Each horizontal row of panels represents a different bin size used for fitting (the bin size can be seen by the spacing of the black points).

Fitting was performed by using a bin-based method. The fitted curves are shown by colorful lines in figure D.13, where different colors correspond to different bin sizes. Figure D.14 summarizes the results. We can see that for all three animal groups there is a range of bin sizes the best fitted values of *k* only very weakly depend on the bin size (for the bin size greater than about 0.3, there are not enough bins to fit the mouse data reliable). The best fitted values fall between approximately 1.5 and 2. In Section 3.3 we assume that *k ≈* 2, see equation (19).

Finally, we will set *k* = 2 and estimate the parameter *λ*, which is related to the ensemble characteristic speed as 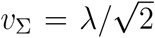, see equation (25). We took

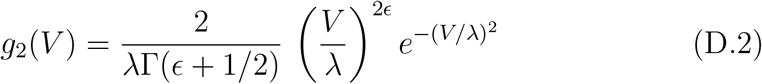

we fitted this function to the empirically obtained distributions (the histograms in figure D.15(a-c)). The best fitting parameter *λ* is shown for different bin sizes in panel (d) of ffgure D.15. The colors correspond to M11 (blue), R13 (yellow), and R15 (green). We observe that the obtained value is only weakly dependent on the bin size (for bin sizes smaller than about 0.3 cm/sec). These values of *λ* (divided by 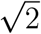) are presented in ffgure 5 with blue symbols; there, for bin-size of 0.215 cm/sec is used.

**Figure C.12:**
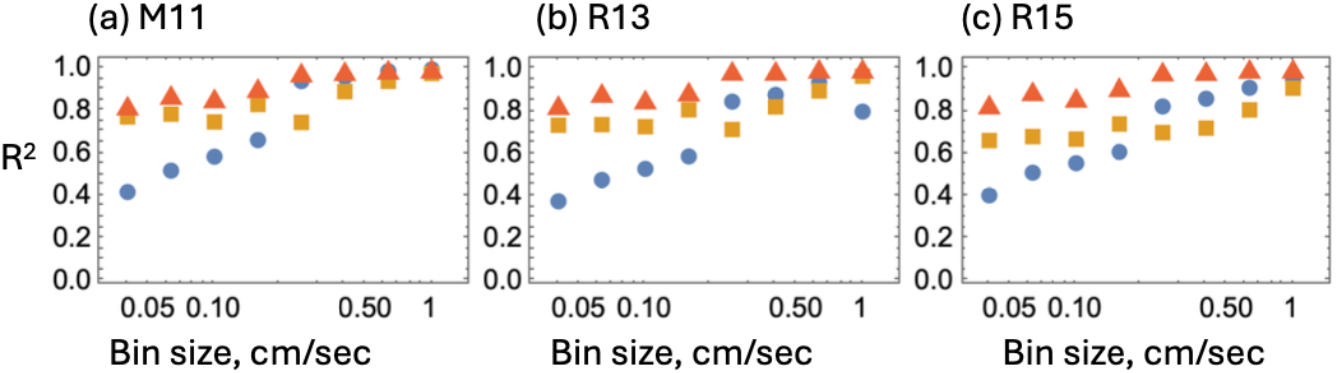
Goodness of fit and the bin dependence of the fitting procedure in figure C.11. The quantity *R*^2^ is shown for two-parametric functions (i) (exponential, red), (ii) (power-law, yellow), and (iii) (Rayleigh, blue), plotted for the fitting procedure corresponding to different bin sizes. A larger value of the *R*^2^ means that the model provides a better fit. (a) M11, (b) R13, (c) R15.

**Figure D.13:**
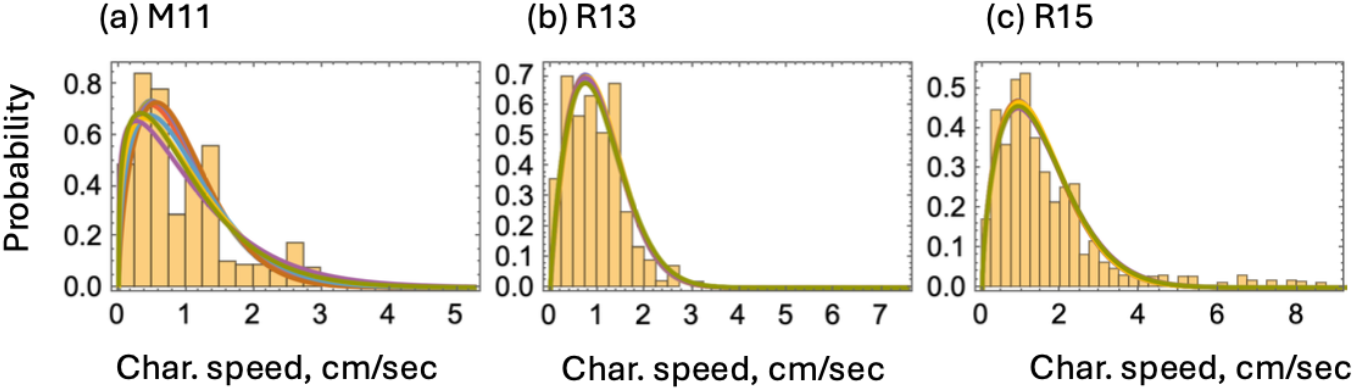
Empirically collected ensemble distributions of characteristic speeds of animals (yellow bars) together with the best fitting functions (equation (D.1), lines). Different colors correspond to different bin sizes from 0.01cm/sec to 0.4 cm/sec. (a) M11, (b) R13, R15.

**Figure D.14:**
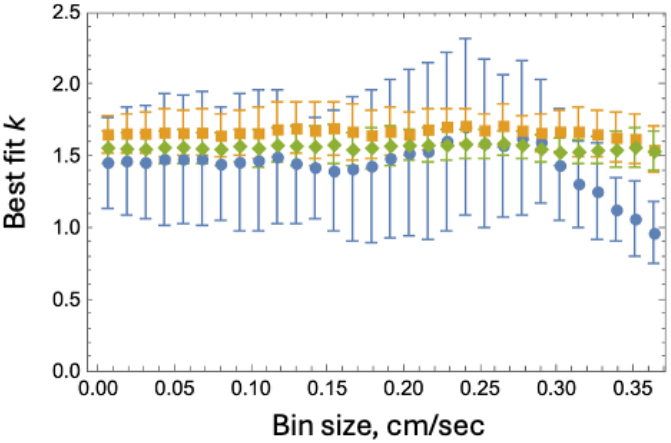
Best fit parameter *k* of the Weibull distribution (equation (D.1)) fitted to empirical ensemble distributions of characteristic speeds, figure D.13. The symbols with error bars represent the best fits with 95% confidence intervals. They are shown as functions of the bin size used, for M11 (blue), R13 (yellow), and R15 (green).

### Derivation of Eq. (8)

We rewrite Eq. (6) using Eq. (7) as:

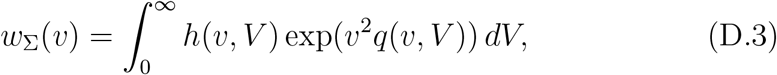

where

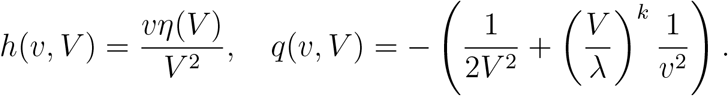

We estimate this integral using Laplace’s method. For ffxed *v* the function *q*(*v, V*) has a maximum at 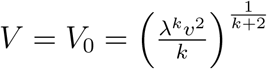. Applying Laplace’s method we have:

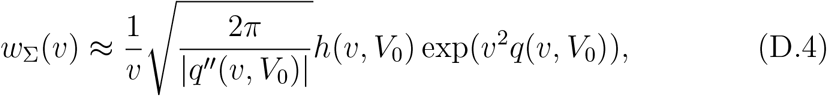

where

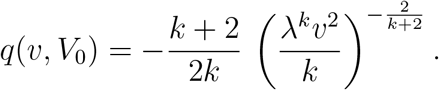

Therefore, the tail of the distribution becomes

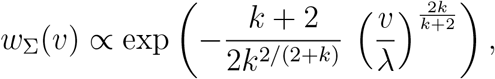

which can be rewritten as

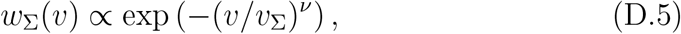

where

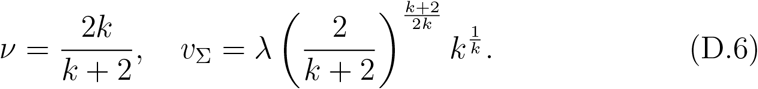

**Figure D.15:**
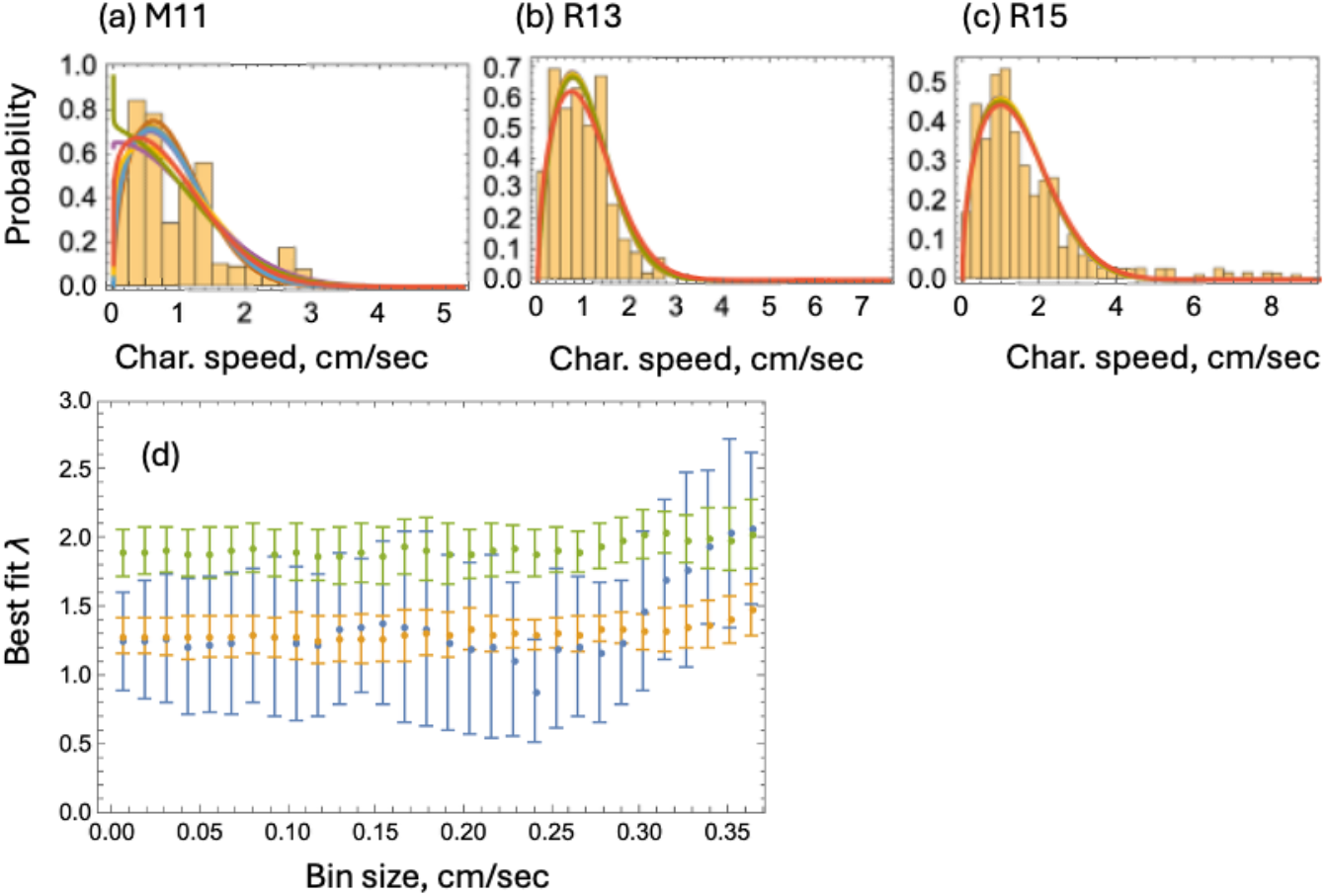
Fitting ensemble distributions of characteristic speeds of animals (yellow bars) with equation (D.2) (lines). (a-c): Different colors correspond to different bin sizes from 0.01cm/sec to 0.4 cm/sec. (a) M11, (b) R13, (c) R15. (d) The best fitted parameter *λ* is shown in as a function of a bin size. M11 (blue), R13 (yellow), R15 (green).

In the special case of *k* = 2 we have *ν* = 1 and 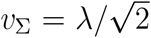. Equation (8) is the normalized tail of distribution D.4, equation (D.5), with *k* = 2.

Note that if the value of *k* is somewhat lower or higher than 2, say between *k* = 1 and *k* = 3 (see ffgure D.14), the tail of this distribution is described by *ν* somewhere between 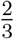 and 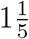, see equation (D.6). We would like to stress that the exponent *ν* does not depend on the choice of *η*(*V*).

